# Evolution of new variants of SARS-COV-2 during the pandemic: mutation limited or selection limited?

**DOI:** 10.1101/2022.09.22.509013

**Authors:** Srashti Bajpai, Milind Watve

## Abstract

The recent pandemic caused by SARS-Cov-2 has witnessed an evolving succession of variants of the virus. While the phenomenon of invasion by immunity evading variants is known for other viruses such as influenza, the dynamics of the ecological and evolutionary process in the succession is little known. Since during the Covid-19 pandemic, large scale epidemiological data were collected and made available in the public domain, it is possible to seek answers to a number of evolutionary questions, which will also have public health implications. We list multiple alternative hypotheses about the origin and invasion of the variants and evaluate them in the light of epidemiological data. Our analysis shows that invasion by novel variants is selection limited and not mutation limited. Further novel variants are not the necessary and sufficient causes of the repeated waves during the pandemic. Rather there is substantial overlap between the conditions leading to a wave and those favoring selection of a partial immune evading variant. This is likely to lead to an association between invasion by new variant and the rise of a new wave. But the association is not sufficiently strong and does not support a causal role of the new variant. The dynamics of interaction between epidemiological processes and selection on viral variants have many public health implications that can guide future policies for effective control of infectious epidemics.

---

The rise of a new variant during an ongoing epidemic of viral infection is an evolutionary process comprising (i) mutational origin of the variant and (ii) natural selection or drift acting on the relative frequency of the mutant. Although both the processes are essential fundamental processes in evolution, one of them can be rate limiting in the given context. If a given mutation offers an all time selective advantage to the mutant, but the probability of the particular mutant arising in the population is small, the rate of evolution will be mutation limited. In contrast, if the population is large enough so that the probability of a specific mutant appearing in the population is quite large, but the conditions required for the selection of the mutant are restrictive and context dependent, then the evolution would be limited by the availability of the selective environment. Careful differentiation between mutation limited versus selection limited evolutionary dynamics can bring about radical changes in perception of a disease and thereby in the public health policy (Vibishan and Watve 2020).

In the field of infectious diseases and public health there are two commonly held perceptions that make this question important. One perception is that mutants that evade host immunity at least partially, are responsible for repeated surges of infection. Such variants may lead to failure of vaccination and the inability to control the epidemic. The second common perception reflected in policies and public information is that rise of new variants can be prevented “…. by reducing the amount of viral transmission and therefore also reducing opportunities for the virus to mutate.” (WHO 2021, University of Meryland Medical System 2021). This perception is based on the underlying assumption that the evolution is mutation limited. Both the perceptions need to be evaluated in order to enhance clarity in perceptions influencing the public health policies.

Many epidemics are known to occur in waves and a number of alternative causes of re-emergence or recurrent wave pattern have been suspected including structured population with heterogeneity in exposure, altered behavior of people and rapidly waning immunity in the population (May and Anderson 1979, Hoe et al 1999, Heffernan and Keeling 2009, Hoen et al 2015, Yang et al 2020). Although multiple possible reasons for the repeated waves in the Covid-19 pandemic are recognized (Epstein et al 2021, Tkachenko et al elife 2021, Cohen et al 2022, Watve and Bhisikar 2021) the predominant popular perception is that new waves were caused by new variants that evaded the immunity conferred by prior infection or vaccination (Thakur et al 2022, Kumar et al 2022, Dutta 2022, El-Shabasy et al 2022, Kupferschmidt 2021). While it has been demonstrated in multiple studies that at least some of the new variants were partially immune evaders (Hayawi et al 2021, Wang et al 2022,) their role as ‘causal’ for new waves is not rigorously examined in comparison with competing hypotheses. Owing to the plausibility of multiple alternative cause effect relationships, which are not mutually exclusive, we need to examine whether during the pandemic, new variants was the only or predominant cause of new waves.

Classically the popular model used for epidemiological predictions is the compartmental or SIR model and its variations. In this model the population is divided into compartments including susceptible, infected and immune or removed. In such a model, immunity is represented as a binary variable. That is an individual is either susceptible or immune. Such models are unsuitable for accounting gradual loss of immunity following natural infections of vaccination. Immunity is a continuous variable in reality and very few models treat it so (Ehrhardt 2019, Watve and Bhisikar 2021). Waves are expected outcomes of the immunity decline models independent of rise of new variants. Therefore loss of immunity and immunity evading variants should be treated as competing but not mutually exclusive causes of recurrent waves and their relative contributions to the waves needs to be evaluated based on their differential testable predictions.

We define below the classes of alternative hypotheses for the evolution of new variants, causal factors for their invasion, beginning of new waves and their interrelationships. We then state the differential testable predictions along with possible falsifying evidence for each of the hypotheses and ultimately test them with public domain data on the Covid-19 pandemic.

## Alternative hypotheses for invasion by new variants and for repeated waves

### A: Invasion by new variants being mutation limited

If the rise of new variants is a mutation limited process then we expect most mutants to arise and invade during the peaks of prior variant waves since mutations are more probable when the standing viral population is high. This is testable by examining the origins of all variants during the pandemic. Furthermore, we can also examine the epidemiological patterns predicted by simulations of a mutation limited model of variant evolution.

Within the mutation limited paradigm there two distinct possible evolutionary paths for new variants.

#### Hypothesis 1: All time selective advantage for new variants with immune evasion

When a prior variant has infected at least a part of the population and the recovered individuals have attained an immune status, any immune evading mutant (all other characters being identical) can be assumed to have a selective advantage. As a result the new variant will start replacing the prior variant at some non-zero positive rate of invasion.

#### Hypothesis 2: Random replacement by drift or selection not related to epidemic parameters

It is possible that new mutants/variants replace the prior one(s) by chance alone such as by genetic drift. It is also possible that there is positive selection on the new variant for reasons that do not affect any of the epidemiological parameters. Intracellular competition, competition for entering a new cell can have trade-offs with net infectivity. Therefore a variant that is a stronger competitor within a host body need not be more infective at an epidemiological level.

### B: Invasion by new variants being selection limited

This school of thought assumes that most of the times during the epidemic the viral population is large enough for ensuring a good probability of an immune evading mutant to arise. Whether the mutant is able to invade the prior variant(s) depends upon the selective conditions present at a given time. Attempts to understand selection on new variants during the pandemic are scanty and we largely lack any useful insights into the nature of selection working on new variants. We propose a descriptive model of how and under what conditions selection is expected to work on immune evading new variants. Further we state the testable predictions of this hypothesis and then tally them with data.

#### Hypothesis 3: The Context dependent selection model

Immune response to a pathogen is multimodal and it is likely that so far we understand only some of the modes of being immune. Immune response is directed to a number of epitopes on the virus and cross immunity across viral variants is well known (Mallajosyula 2021). Mutations do not alter all of the epitopes together and therefore immune evasion is always partial. Although it is difficult to decide the exact level of immunity required to give protection from an infection, it is generally agreed that the titers achieved after a natural course of infection or vaccination are many fold higher than the minimum required for immunity. As a result, immunity following natural infection or vaccination is most likely to be effective against the partially immune evasive variants as well (Zeng et al 2022, Tragoning et al 2021). Therefore, when immune levels are high, partial immune evaders would be unable to invade. Perhaps for the same reason, even after omicron was the predominant veriant, omicron specific vaccines were not found to give greater protection than prior vaccine boosters (Callaway nature 2022). Variants are unlikely to have any selective advantage when the hosts are fully immune as following vaccination or natural infection. Acquired immunity is known to decline with time with varying rates. The decline appears to be considerable in Covid-19 (Dolgin 2021, Feikin et al 2022). Models involving declining immunity show that if the population immunity declines below a threshold, a new wave is likely even when there is no new variant (Ehrhardt 2019, Watve and Bhisikar 2021). However, the stage just prior to this offers the right opportunity for a new variant with partial immune evasion to get selected. Because of partial immune evasion the threshold for the beginning of new wave for the new variant is likely to be reached earlier than that for the prior variant. As a result, just before a new wave begins, or in the early phase of a new wave a new variant is most likely to invade and replace the prior variant(s). Such a process of conditional selection of immune evasive mutant will still lead to an association of new variant with new wave, but the cause effect relationship may not be simple.

Conditions that lead to a new wave are also the conditions favorable for a new variant. Therefore although the new variant may not be causal to the wave, it will ride the wave. The epidemiological predictions of this model are different than the model in which a new variant “causes” the new wave.

Although we used immunity decline as a variant independent cause of repeated waves, other causes for the wave pattern have been suggested such as cyclic changes in the population behavior (Tkachenko et al elife 2021). Our model can conceptually incorporate other causes of wave pattern without much change in the model structure. Although multiple reasons can be responsible for the wave pattern, as long as conditions triggering a new wave overlap with conditions favoring selection of a new variant, the model predictions remain the same.

## Differentiating between the alternative hypotheses

By hypothesis 1, the majority of invasions should begin close to the peak of incidence of prior variant. If the new variant has a higher rate of transmission, the invasion should be accompanied by an increase in slope of the incidence curve. If the rise in slope is sufficiently large a detectable new wave can be created. The new wave is expected to be characterized by a continued or declining slope of the prior variant and the rise in slope of the incidence curve being entirely attributable to the new variant (fig 1a).

**Figure 1:**
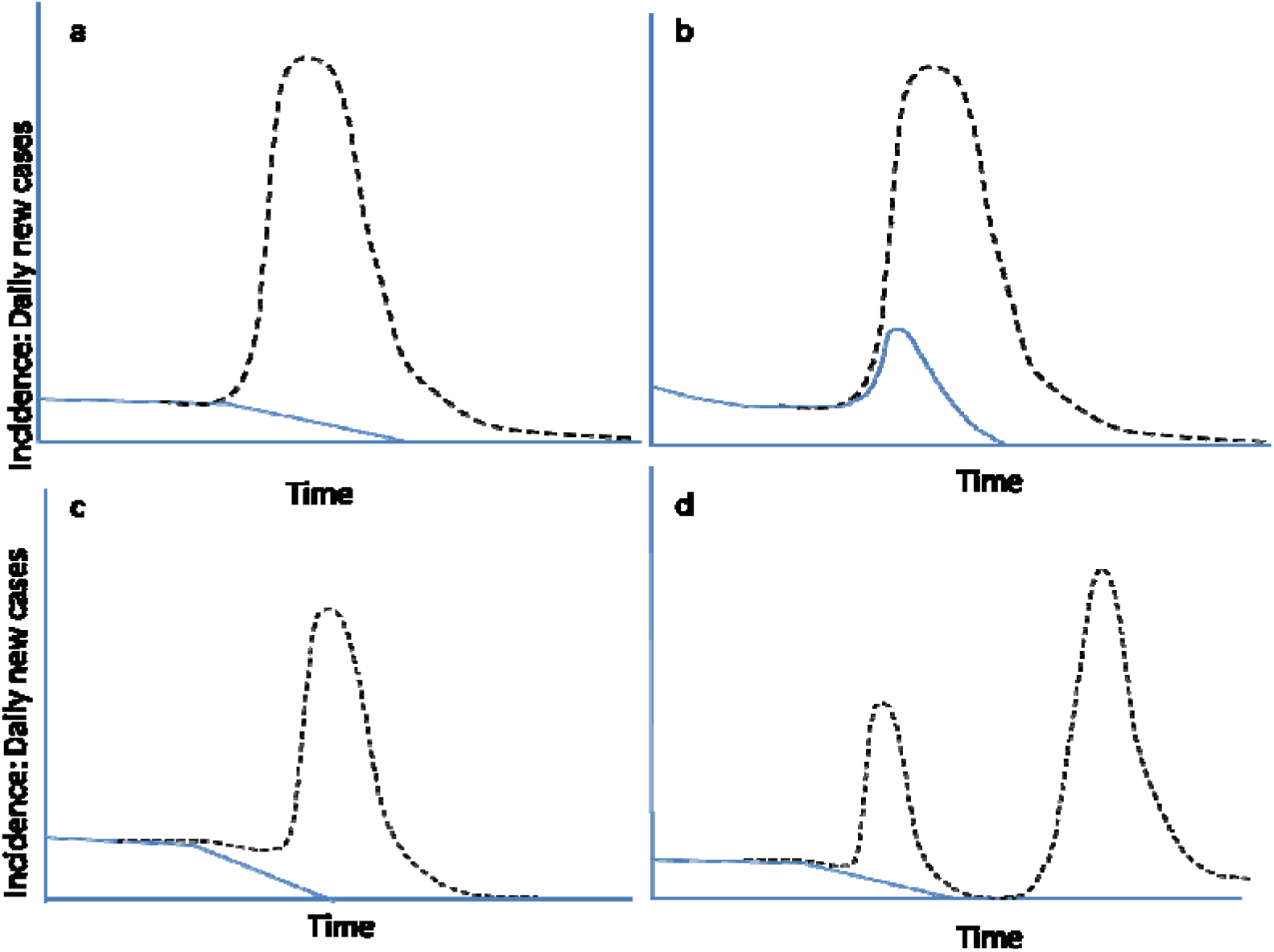
Solid line indicates incidence by the prior variant(s), dashed line the total incidence (a) A hypothetical expected incidence curve if a new variant is the only necessary and sufficient cause of the wave.. We can visualize four different ways in which the relationship between the total incidence and proportion of prior and new variants differs from the expectation of hypothesis 1. (b) Prior variant wave: In this pattern, the incidence trend of the prior variant(s) also shows an upward shift during early phase of the wave i.e. even if we remove the new variant, there is a detectable wave. (c) Prior replacement: A partial or complete replacement of old variant(s) by a new one happens before the wave begins. (d) Complete wave course without a new variant: A wave arising and falling i.e. completing its course with no new variant arising during this course. This can happen if a new variant has completely replaced the prior one(s) before the beginning of a new wave. This pattern indicates that a new variant is neither necessary nor sufficient for a wave.

By hypothesis 2 the probability of new variant invading will either be directly proportional to standing viral population, or constant in time. The testable prediction of this hypothesis is that the probability of new variant arising in a given time interval will be proportionate to the area under the incidence curve or constant over time depending upon whether selection unrelated to epidemiological parameters or drift is the predominant factor.

By hypothesis 3 when a new wave begins, the incidence by prior variant(s) is also likely to go up at least in the initial phase of the wave (fig 1b). Alternatively the new variant need not necessarily have a higher rate of transmission therefore it might replace the prior variant(s) without causing a wave (fig 1c). It is also possible that new waves arise without any detectable invasion by a new variant (fig 1d). This hypothesis also expects that new variants are unlikely to invade at or just after the peak of prior wave when host population immunity is at its maximum. Invasion by new variants can happen only after substantial time gap following the peak of prior wave when population immunity has sufficiently declined to allow a new surge.

If the new variant is more infective than the prior one(s) at any given phase of the epidemic, we would expect the slope of the total incidence curve to increase as a function of the standing proportion of the new variant or to the rise in proportion of the new variant. In the short run it can be assumed that the host immunity status may not have changed sufficiently to affect the correlation. In the long run as the host population acquires immunity to the new variant, the correlation may change. Therefore such correlations should be sought only over a short time span before the new variant proportion saturates and/or the wave reaches its peak, whatever is earlier.

We also use some simulations here as a tool with limited implications. Using simulations we test the notion qualitatively that if mutations are limiting and an immune evading mutant has an all time advantage, then most invasions will begin near to the prior peak. Since we do not have empirical information of many parameters, simulations cannot be used to make any quantitatively useful predictions.

## Methods

### Simulations

Since most conventional models are based on the compartmental SIR models, we start with a net population of unity and *S(0), I(0)* and *R(0)* the initial fractions of susceptible, infected and immune individuals respectively. With a starting variant *V_0_* the SIR model is run with a probability of *S* to *I* conversion = *K_1_.S.I*, the probability of *I* to *R* conversion = *K_2_.I*. The probability that a new variant *V_1_* (and thereafter serially to V_n_) is generated at a given time is directly proportionate to *I*. A new variant is assumed to be generated when a randomly drawn number is < *p.I* and *V_n_(t)* takes a small value of *0.001*. For the new variant *S_n_(t)* = *1− I(t)* and *R_n_* = *0* and the simulation equations apply to the variant in a similar way. These simulations assume that immunity to each variant is acquired independent of other variants and there is no cross immunity. Since in typical SIR models, immunity is treated as a binary variable, it is difficult to incorporate cross immunity or gradual loss of individual immunity.

There are models that incorporate individual immunity as a continuous variable. We use the Watve and Bhisikar (2021) model, which is able to predict repeated waves in the absence of any immune evading variant. We compare and contrast the predictions of the two models so that we have some simple qualitative differential testable predictions for the hypotheses of our interest. We do not use the simulations to make any quantitative predictions.

#### Source of data

Data for testing the predictions were obtained from the public domain database https://ourworldindata.org/explorers/coronavirus-data-explorer (Ritchie et al 2022). The database has daily details of country specific registered cases and deaths. In selected countries having adequate sequencing efforts there is bi-weekly updated proportion of recognized variants of concern among the ones sequenced. The limitations of the data are set by the data recording accuracy that might differ across countries. Since only a small fraction of the samples are processes for identification of variants we can talk about when and where a variant was first detected, which is not necessarily the same as when and where a variant originated, but a reasonable reflection of it. The samples processed for sequencing are not randomized and some selection bias in favor of new variants is likely since suspected new variant cases are more likely to get sequenced. Also there can be variable delays in sequencing. Owing to the data limitations the following specifications and working definition are used in the analysis. We consider how the limitations and biases may affect the inferences in the sensitivity analysis section later.

#### Choice of Variants for the analysis

We use the variants being monitored (VBM) as listed by CDC in https://www.cdc.gov/coronavirus/2019-ncov/variants/variant-classifications.html#anchor_1632158885160, namely the ones with respective WHO labels alpha, beta, gamma, delta, epsilon, eta, iota, kappa, zeta, mu and omicron and its sub-variants. Data on all the variants are used for analyzing predictions related to the origin of variants. However since delta and omicron have been largely blamed for the second/third waves and subsequent waves respectively, we use these two variants for analysis related to the association of variants with specific waves.

A data segment is taken as two weeks since the variants data available are proportions among sequenced samples over two weeks.

#### Origin of a variant

The working definition for the origin of variants is the date and country where the variant was first detected.

#### Invasion by a variant

For each of the countries having variant data, the date from which the proportion of a variant among the sequenced genomes starts monotonically increasing at least for three consecutive data segments is taken as the working definition of beginning of invasion by the variant.

The working definition of a wave is one in which a rise in the daily registered new cases is at least by a thousand from the baseline. A peak is recognized by monotonic increase for at least two data segments followed by monotonic decrease for at least two data segments.

A data interval used for studying correlation between proportion of variant and rate of transmission: the data interval used begins when the new variant invasion begins or a new wave begins (whichever is earlier) and ends when the proportion of new variant saturates (most often near 100%) or the peak of the wave is reached, whichever is earlier. The association analysis is restricted to this interval because beyond that interference by host immunity to the new variant may change and even invert the correlation.

## Results

In the absence of empirical estimates of many of the parameters, the simulations cannot be used to predict any quantitative patterns. They have a limited use to predict some contrasting qualitative patterns. Assuming mutation limited evolution of variants and new variants as the necessary and sufficient cause of waves, we see that most invasions begin near the peaks of prior variant incidence and the total incidence always remain high (fig 2a). There are stochastic fluctuations but a significant time gap with low incidence level is not observed. In contrast the Watve and Bhisikar (2021) model treating immunity as a continuous variable, is able to predict a realistic life history of a wave including a low incidence endemic like state between two waves.

**Figure 2:**
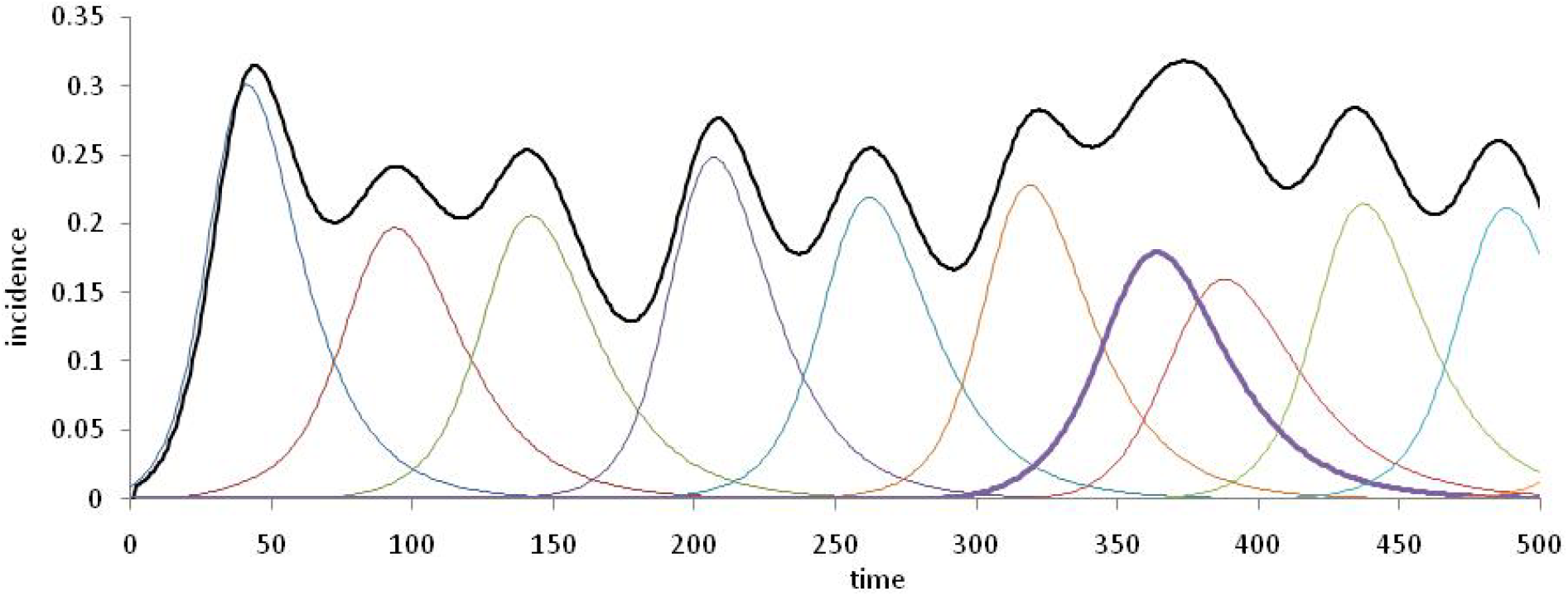
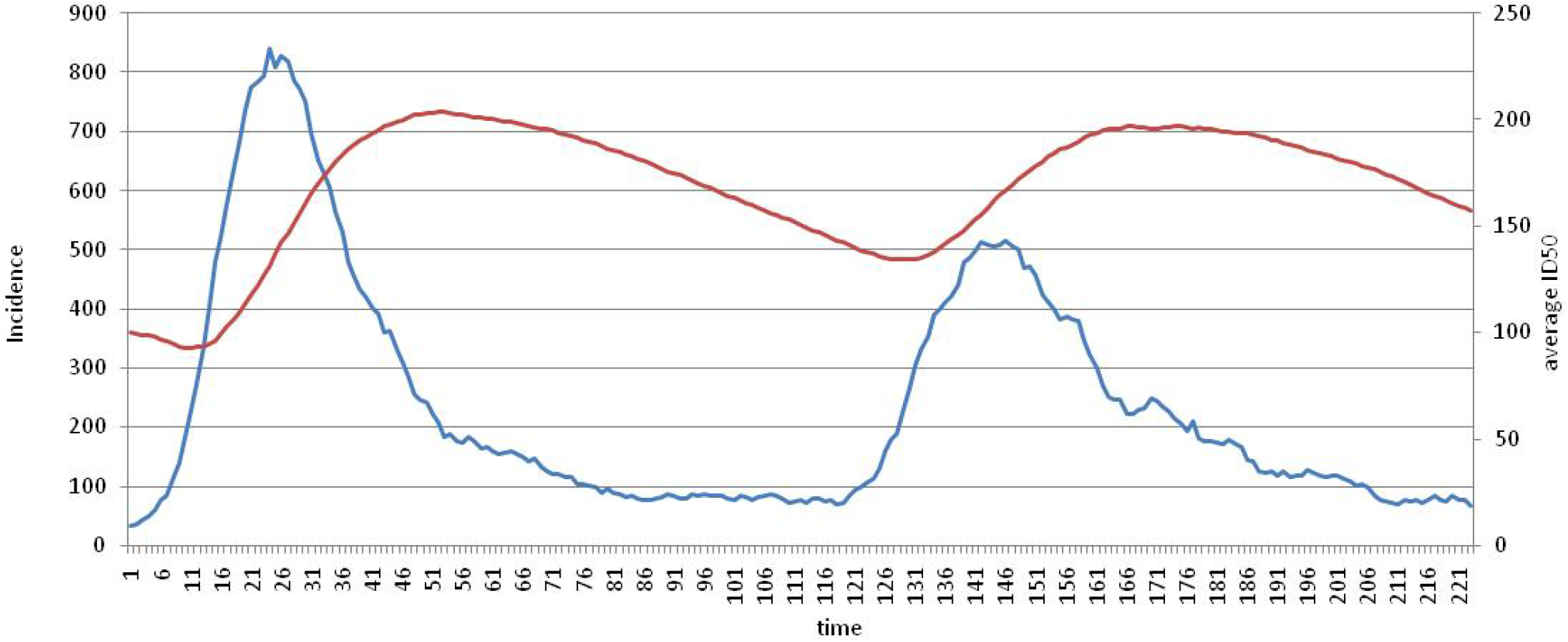
Example patterns seen in simulations using the two contrasting models. a. SIR model incorporating generation of novel immune evading random mutations. The black line represents total incidence and all coloured lines the incidence by the succession of variants. Note that most of the new variants invade near the peaks of prior variants since the probability of mutation is highest that time. b. A result of the Watve and Bhisikar (2021) model in which waves are separated by a considerable period of low incidence apparently endemic state. The blue line represents in the incidence curve and the red line the mean population immunity. New wave can be triggered when the population immunity declines below a threshold. In this model, waves are seen in the absence of a new variant.

For testing the predictions related to the association between new variant and new wave, adequate data on variants were available for 125 waves from 64 countries. We observe that out of the 125 waves not a single wave complies to the expectation of hypothesis 1 as in fig 1a, 72 waves show prior variant wave as predicted in fig 1b, in 92 there is prior invasion of new variant much before the beginning of the wave as in fig 1c and 27 waves complete their course without the appearance of any new variant. Figure 3 shows representative patterns corresponding to fig 1b to 1d from one country each. All the 125 waves have at least one of the three patterns contradicting hypothesis 1.

**Figure 3:**
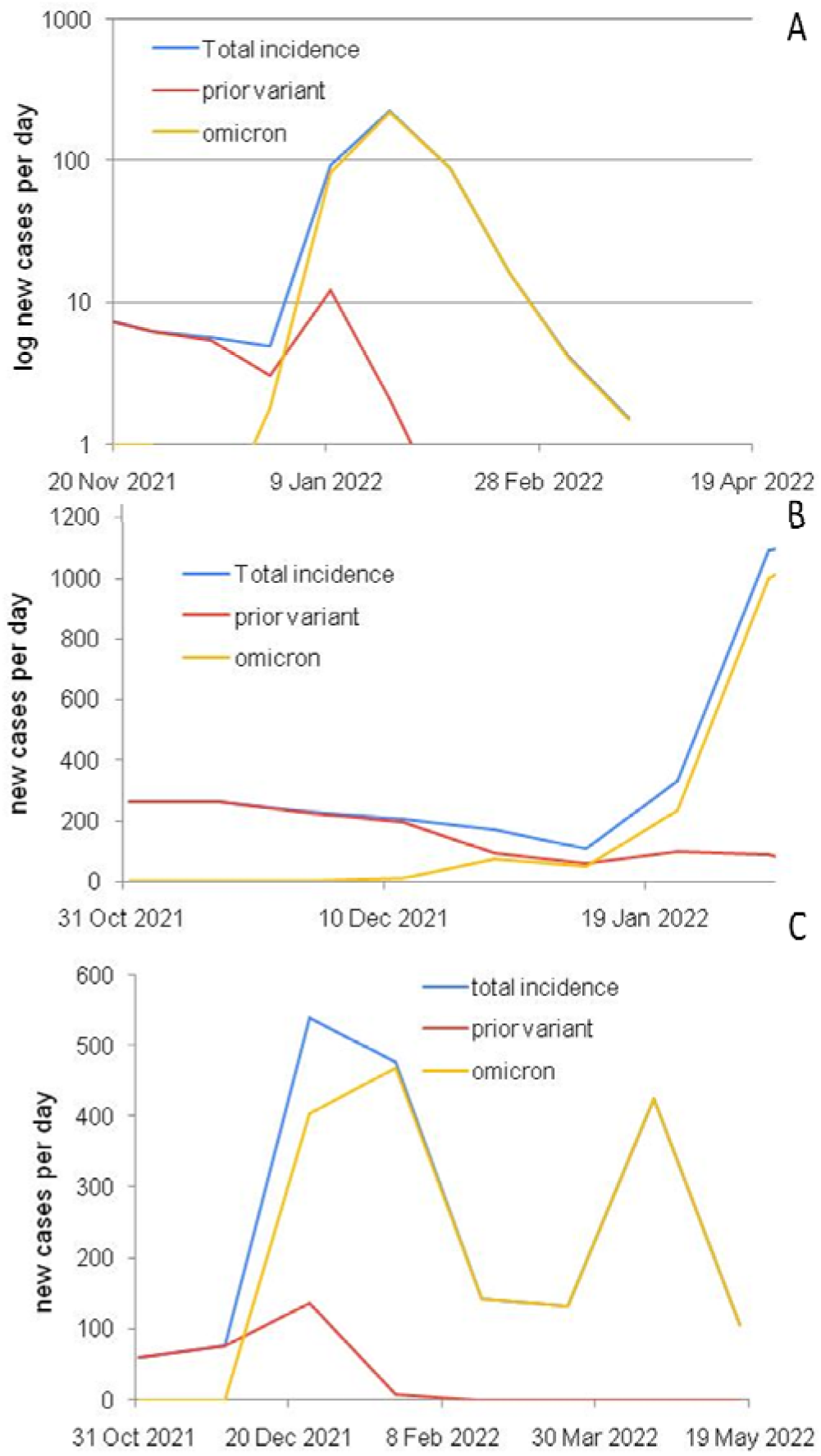
Sample incidence curves showing deviations from the pattern expected by hypothesis 1. a. Prior variant wave: the incidence of infection by the prior variants also increases in the first phase of the wave. Data shown from omicron associated wave in India. b. New variant prior invasion: Invasion by a new variant begins and old variants are replaced to a substantial extent much before the beginning of the new wave. Data from Russia at the beginning of the omicron associated wave. Note that omicron replaced the prior variant delta by about 50% while the total incidence was decreasing. c. A wave without any new variant: The second wave seen in this figure from Canada was apparently without a new variant.

Further, the correlation between standing proportion of new variant or increase in the proportion of the new variant is poorly correlated with the increase in slope of the incidence curve.

Correlations for both the delta and omicron are weak and not even 1 % variance in the slope of incidence curve is explained by a new variant (fig 4 a,b,c,d). The correlations do not support the notion that the increasing slope at the beginning of the wave is because of the new variant being more infective. Thus not only the prediction of hypothesis 1 is not found to be true, it can be said to be consistently rejected by multiple prediction tests.

**Figure 4:**
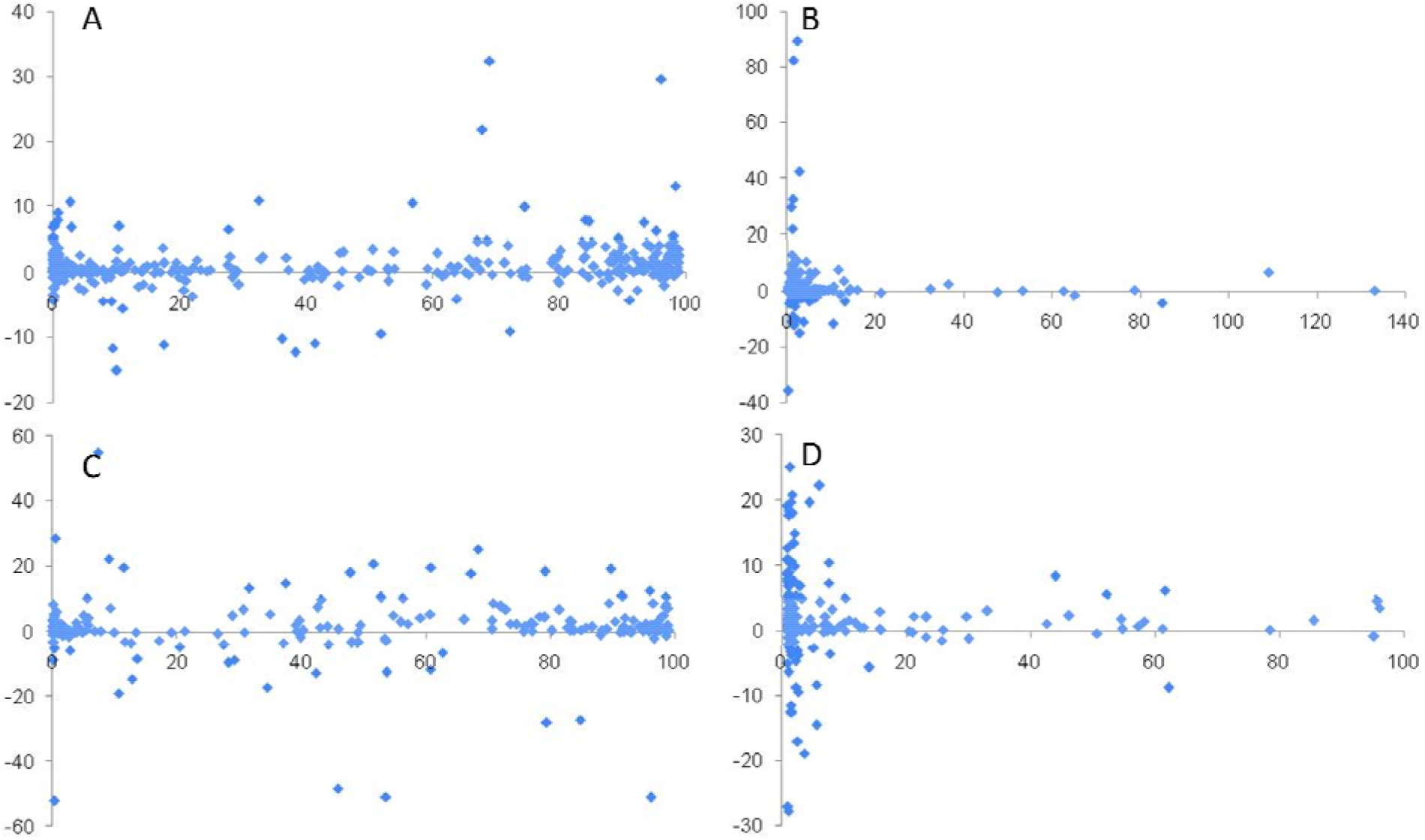
Correlations between the standing proportion of the new variant and change in slope of incidence curve (left panel) and between proportionate increase in the new variant and change in slope of incidence curve (right panel). A and B: data for delta variant associated waves, C and D for omicron associated waves.

Plotting the origin of variants along with the incidence curves shows that unlike the general prediction of the mutation limited hypotheses, we see that most of the variants have originated at low levels of incidence (fig 5). Since there are only 16 variants or subvariants with distinct identifiable origins, we did not perform a quantitative analysis for whether the probability of a new variant arising is constant in time or proportionate to the area under the curve, but since 14 out of the 16 origins are at lower levels of incidence, the data does not resonate with prediction of the mutation limitation paradigm.

**Figure 5:**
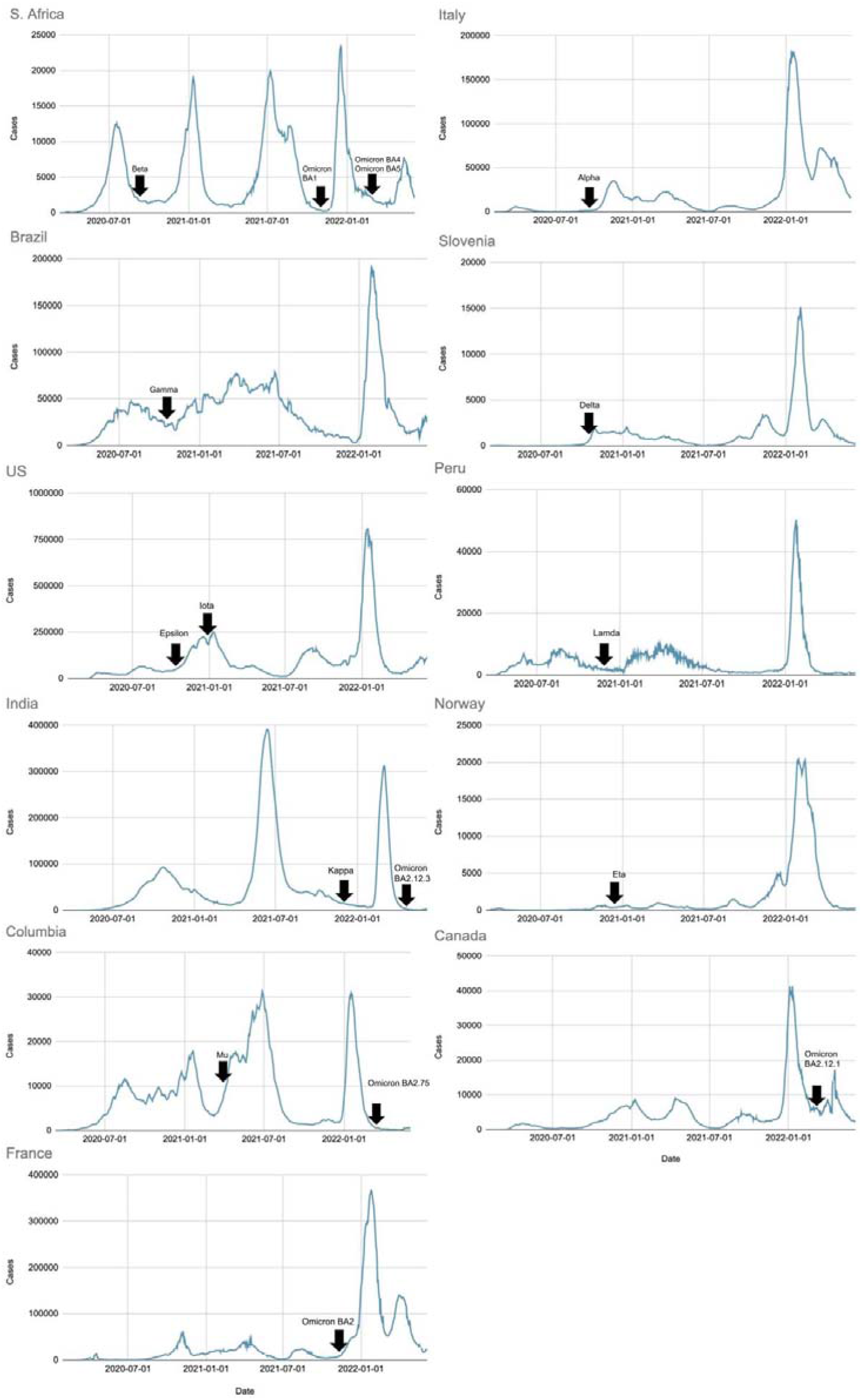
The countries and time of origin of the 16 variants and sub-variants being monitored. Origin or a new variant is shown by an arrow along the incidence curve. Note that most variants are first detected when the incidence was low.

Further if we look at the beginning of invasion by new variants across all countries that have variant data, the invasions also begin most commonly when the incidence is low. It is neither distributed randomly in time nor proportionate to the area under the curve rejecting drift or independent selection hypothesis. On the contrary the beginning of invasion is most commonly placed after a certain time gap from a prior peak (fig 6) as expected by hypothesis 3.

**Figure 6:**
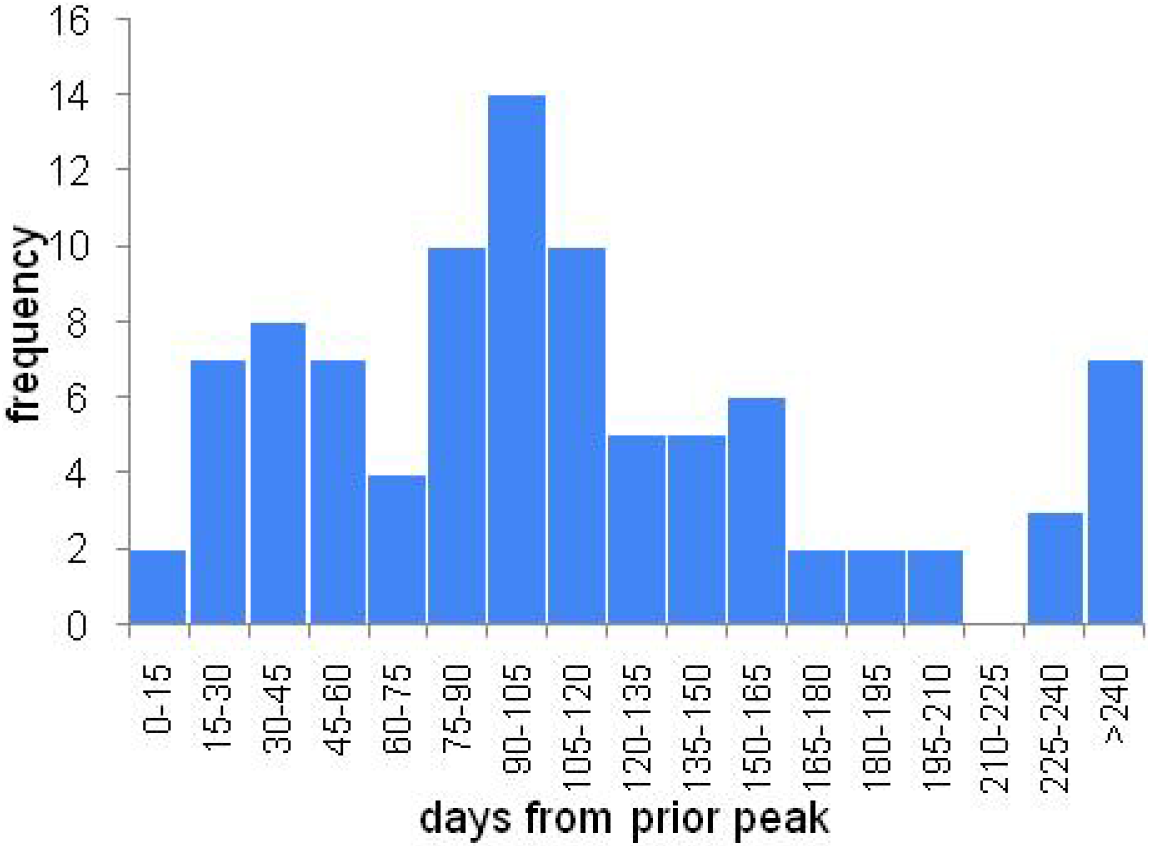
The frequency distribution of the gap between prior peak and new variant invasion. The mode is at about 100 days. Note that very few invasions happen just after the peak unlike the expectation of the mutation limited paradigm.

### Sensitivity analysis

We examine now whether the patterns observed could be the result of the unavoidable biases and limitations in the data. The variants are defined as lineages and data are available only on the variants of concern defined by WHO. However, mutations keep on accumulating within the variants and sub-variants. Not all have been named and thereby data on their frequencies unavailable. Therefore the presence of waves without new variants (fig 1 d and 2c) cannot be said to be a strong falsifying evidence against the hypothesis 1. It is possible that the waves are associated with some mutations about which we do not have data. Therefore we avoid over-interpreting the pattern in figure 1d and 2c, i.e. waves without new variants. The apparent time lags between rise or immigration of a new variant and its first detection can bias the time of origin or time of invasion. The actual origin or beginning of invasion would be earlier than what appears in the data. Any correction for this bias will actually make prior invasion pattern (fig 1 c and 2b) stronger than what is apparent. Therefore this is a strong reason to reject hypothesis 1. The bias in the samples selected for sequencing is likely to lead to overrepresentation of new variants as admitted by (Ritchie 2020). In spite of this bias we see prior variant waves (fig 1b and 2a) in 72 cases. Any correction for this bias would actually strength the prior variant wave pattern. Therefore this can also be treated as a strong evidence for rejecting the mutation limited perspective. The data intervals selected for the correlation analysis avoid ranges in which biases would arise. For example we avoid taking the slopes in the receding part of the wave which can be said to be due to the population gaining substantial immunity to the new variant. We also avoid taking data after a variant completely replaces the prior variant or reaches saturation since the subsequent flattening of the curve would weaken any correlation. Therefore the overall correlation between rise in proportion of new variant and rise in the apparent rate of transmission can be said to be inherently weak with confidence. Therefore although data bias may make some of the evidence weak, there are still substantial grounds to reject hypothesis 1. The selection limitation paradigm, assuming immunity decline to be a common cause for a new wave as well as for selection of a partial immune evading variant, gains support throughout the analysis.

## Discussion

The epidemiological patterns in the origin and invasion by new variants and origin and rise of new waves in the incidence curves contradict the mutation limited paradigm and support the selection limited paradigm. The immunity decline as the common cause for rise of new wave as well as selection of new variants looks more promising. This may have a broader application covering influenza, common cold and other viral infections in which new variants arise very frequently. Owing to better collection of data in the public domain, testing multiple differential predictions was possible in the Covid-19 pandemic, in the light of which revisiting influenza may be revealing. The question why variants arise frequently in influenza and corona viruses but not in Pox viruses or poliovirus for example remains open and the answer may lie in the dynamics of immunity. Mutation rates are unlikely to be very different across viruses. If the rate of immunity decline is rapid, the frequency or new variants appearing should be high by our hypothesis. Immunity to pox virus or polio virus is known to be long lasting and that might arrest the evolution of new variants. Such new possibilities suggested by this analysis need serious exploration theoretically as well as epidemiologically.

The selection limited evolution also necessitates a substantial revision of the mainstream thinking about the epidemic and the mitigation measures. Trying to limit the transmission by lockdown measures to arrest new variants as advocated by WHO and other health authorities (WHO 2021, University of Meryland Medical System 2021) is unlikely to work because of multiple reasons. First the lockdown measures have not been shown to be very effective in curbing the transmission except for the very initial phase of the epidemic (Kharate and Watve 2022, Yanovskiy and Sokol 2022, Herby et al 2022). Secondly rise of new variants not being mutation limited, limiting the viral population in realistic limits may not prevent rise of new variants. Viral populations even within one host individuals are quite large and so are mutation rates in RNA viruses (Carrasco-Hernande et al 2017). Therefore mutants are expected to arise very frequently. Whether conditions favor selection of these mutants should be the more relevant question shaping the evolution of the virus. The declining immunity driven selection has the potential to give more useful insights into viral evolution and needs to be pursued more seriously at theoretical as well as empirical level.

On this background the relevant questions are whether we can shape the selective conditions for new variants by appropriate public health strategies. How would the alternative non-pharma preventive measures alter the selective landscape, how vaccines and boosters would shape the selective landscapes are open questions inviting theoretical as well as empirical work. An insightful understanding of the selection acting on the virus variants is likely to refine and appropriately design preventive strategies with long term effects in future epidemics.

## Notes

https://ourworldindata.org/explorers/coronavirus-data-explorer

### Competing Interest Statement

The authors have declared no competing interest.

